# The cost of stressing the royal family: parental glucocorticoid exposure has delayed adverse effects on chick physiology and survival in king penguins

**DOI:** 10.64898/2026.11.16.767074

**Authors:** Anais Cotton, Frédéric Angelier, Sandra Avril, Lucie Abolivier, Jean-Patrice Robin, Pierre Bize, Vincent A. Viblanc, Pierrick Blanchard, Antoine Stier

## Abstract

When exposed to stressful environmental conditions, adults of long-lived species may favour self-maintenance at the expense of reproductive investment. In breeding king penguins (*Aptenodytes patagonicus*), we previously showed that prolonged glucocorticoid hormone exposure during early breeding resulted in surprisingly little effect on parental self-maintenance. Here, we investigate whether the potential costs of this experimental stressor could be transferred to offspring through altered parental care, negatively affecting their growth, physiology and survival. We exposed parents to a ca. 1-month elevation in corticosterone and monitored parental investment and offspring performance over one year, from incubation to fledging. Corticosterone-treated parents exhibited reduced prolactin levels and spent less time at sea, highlighting alterations in both endocrine and behavioural processes supporting parental investment. Yet, parental care behaviour, hatching success and chick body mass/size were not significantly impacted. Major delayed costs of the parental treatment were however evident, as chicks from corticosterone-treated parents exhibited accelerated postnatal telomere shortening and faced nearly twice the mortality hazard of chicks from placebo-treated parents. By revealing the hidden reproductive costs of apparent parental resilience, our study challenges how we assess the consequences of environmental stressors in long-lived species, particularly when their effects only emerge long after exposure.

## Introduction

Prioritising self-maintenance over current reproduction may help parents cope with environmental challenges and preserve their survival and future reproductive opportunities (Wingfield & Sapolsky, 2003; Lemaître et al., 2015). However, this allocation shift can impose both short-term costs on offspring, including lower growth and short-term survival (Monaghan, 2008; Badyaev & Uller, 2009; Spée et al., 2011; Kidawa et al., 2017; Pegan et al., 2019), as well as long-term costs revealed for instance by reduced telomere length (Herborn et al., 2014; Pegan et al., 2019; Casagrande et al., 2020), a molecular biomarker of ageing and predictor of lifespan and lifetime reproductive success in birds (e.g. Eastwood et al., 2019; Heidinger et al., 2021; Chik et al., 2024; Novelli et al., 2026). To regulate this balance, hormones play a central role: glucocorticoids coordinate physiological responses to stressors and may favour self-maintenance (Wingfield & Sapolsky, 2003), whereas prolactin supports parental behaviours such as incubation, brooding, and chick provisioning in birds (Angelier & Chastel, 2009).

Rather than exerting consistently negative or positive effects on reproduction, glucocorticoids and prolactin can influence parental investment and offspring performance in context-dependent ways, shaped by species-specific life histories, environmental conditions and individual characteristics (Angelier & Chastel, 2009; Crossin et al., 2015). For instance, elevated parental glucocorticoids increased maternal foraging effort and chick mass in macaroni penguins (*Eudyptes chrysolophus*; Crossin et al., 2012), whereas the same treatment reduced reproductive output by 42% in Adélie penguins (*Pygoscelis adeliae*), without affecting the mass of surviving chicks (Thierry et al., 2013a). Similarly, experimentally reduced prolactin levels decreased incubation commitment and hatching success in Adélie penguins (Thierry et al., 2013b). Consistently prolactin remained elevated during prolonged parental absences at sea in emperor penguins (*Aptenodytes forsteri*), potentially supporting continued parental care (Lormée et al., 1999). Environmental and life-history contexts also modulated glucocorticoid effects in black-legged kittiwakes (*Rissa tridactyla*), with elevated parental corticosterone reducing chick growth and reproductive success in Pacific populations but producing opposite responses in Atlantic populations (Schultner et al., 2013). Finally, elevated corticosterone during chick rearing was associated with higher reproductive success but shorter adult telomeres in male common terns (*Sterna hirundo*), suggesting potential physiological costs of increased reproductive investment (Bauch et al., 2016).

Species regularly exposed to predictable environmental challenges may have evolved mechanisms that limit the reproductive consequences of hormonal changes, allowing them to maintain reproductive investment (Lynn & Benowitz-Fredericks, 2024). Nevertheless, reducing reproductive investment may remain advantageous when environmental challenges become severe or unpredictable, favouring flexibility over consistently maintained reproductive effort (Wingfield et al., 1998; Wingfield & Sapolsky, 2003). Whether such flexibility can be maintained under prolonged glucocorticoid elevation, and at what cost to subsequent reproductive stages and offspring development, remains poorly understood in the wild. Seabirds provide a relevant model for investigating the reproductive consequences of sustained endocrine changes, given their high adult survival, delayed maturity, low annual reproductive output and recurrent exposure to environmental challenges (Weimerskirch et al., 1992; Hamer et al., 2001). Their strong reliance on adult survival and residual reproductive value suggests that self-maintenance should be favoured over current reproductive investment under stressful conditions (Culina et al., 2019). Among them, king penguins (*Aptenodytes patagonicus*) provide an ideal model for investigating how glucocorticoid elevation during early breeding affects parental investment and long-term offspring performance, since their reproductive cycle lasts more than one year (Weimerskirch et al., 1992). This allows assessment of immediate effects and delayed costs after the stressor. Previous correlative work in this species showed that individuals abandoning reproduction following severe depletion of energy reserves exhibited elevated corticosterone (CORT), the primary avian glucocorticoid, and reduced prolactin levels (Groscolas et al., 2008). More recently, using the same experimental manipulation as in the present study, we showed that prolonged CORT elevation during early breeding induced relatively limited changes in adult physiological traits, most of which were transient or small in magnitude (Cotton et al., 2026). These findings suggest some physiological buffering against sustained glucocorticoid elevation, but whether this apparent resistance also extends to reproductive investment, or instead involves costs to offspring development and survival, remains unknown.

In this study, we experimentally elevated CORT in free-ranging king penguins during early breeding. Breeding adults received either a corticosterone-releasing implant (CORT-treated group) or a placebo implant (Placebo group), resulting in elevated corticosterone levels for approximately one month in treated individuals (Cotton et al., 2026). We subsequently assessed (1) endocrine and behavioural traits associated with parental investment, including prolactin levels, time spent at sea, and chick care behaviour, and (2) offspring performance, including hatching success, chick body mass and size growth, as well as relative telomere length and survival until fledging. We considered two alternative hypotheses. First, if king penguin parents can buffer both their own physiological maintenance (Stier et al., 2019; Cotton et al., 2026) and reproduction while facing prolonged glucocorticoid elevation, CORT-treated parents are expected to maintain parental investment and offspring performance at levels comparable to Placebo. Alternatively, if prolonged glucocorticoid elevation favoured parental self-maintenance at the expense of current reproduction, CORT-treated parents are expected to exhibit reduced prolactin levels and altered parental behaviours, potentially resulting in lower hatching success, reduced chick growth, accelerated telomere shortening and decreased offspring survival.

## Materials and methods

This study was conducted during the 2018-2019 breeding season at La Baie du Marin, Possession Island, Crozet Archipelago. King penguins alternate incubation and chick-guarding duties with foraging trips at sea. After about 53 days of incubation, chicks remain under continuous parental attendance during the guard phase until they become thermally independent at about 35 days of age (Duchamp et al., 2002).

A total of 49 breeding pairs were monitored from courtship in early November through the end of the guard phase (the same individuals as in Cotton et al. 2026). Breeding pairs were randomly assigned to female-treated (n = 23) or male-treated (n = 26) groups, with only one parent per pair receiving an implant. Within each group, adults received either a subcutaneous corticosterone-releasing implant (CORT; NG-111, 50 mg, 90-day release; 8 × 3 mm) or an identical hormone-free placebo implant (NC-111; Innovative Research of America, Sarasota, FL, USA). Among female-treated pairs, 12 females received CORT implants and 11 received Placebo implants; among male-treated pairs, 13 males received CORT implants and 13 received Placebo implants.

At D0, blood from adults was sampled immediately before implantation, which occurred during the first incubation shift in females and the second incubation shift in males. In both sexes, implantation occurred three days after returning from their first foraging trip following egg laying, standardising fasting duration between sexes. Additional blood samples were taken six days after implantation, and after the first and second post-implantation trips at sea (see details in Cotton et al., 2026). Daily observations of marked individuals enabled to assess the time spent at sea, hatching date and thus to dissociate two breeding stages: incubation and chick-guarding. Parental chick-care behaviours (i.e. chick grooming, feeding or other direct interaction) were monitored using scan-sampling throughout the guard phase (i.e. 283 ± 120 scans per individual, mean ± SD; n = 41 individuals). Plasma prolactin concentrations were measured at D6, L1, and L2 using a radioimmunoassay following Spée et al., (2010). Intra- and inter-assay coefficients of variation were 8.93% and 11.27%, respectively.

### Chick monitoring and measurements

Chick mass, total bill length and mean flipper length were measured at 10, 20, 35, 100, and 150 days post-hatching, as well as at the end of the pre-fledging moult (approximately 300 days). Structural size was assessed using the first principal component (PC1) from a principal component analysis (PCA) of standardised total bill and mean flipper lengths. Measurements were pooled into a single PCA to obtain a common structural size index across development, with PC1 explaining 98.3% of the total variance. Chick survival was checked daily during the guard phase and then twice a week until fledging.

Blood samples collected at 10 and 35 days post-hatching, the two sampling ages with the largest sample sizes, were used for molecular sex determination and measurement of relative telomere length (rTL). We quantified rTL by quantitative PCR (qPCR) as the ratio of telomeric repeat abundance to single-copy reference gene abundance (T/S ratio), following established protocols (Stier et al., 2014; Cotton et al., 2026). Technical repeatability, assessed by measuring the same samples across qPCR plates, was 0.88 ± 0.03 SE (95% CI: 0.82–0.93; LRT *p* < 0.001). Within-individual biological repeatability of rTL between days 10 and 35, with paired samples analysed on the same plate, was 0.68 ± 0.10 SE (95% CI: 0.44–0.82; LRT *p* < 0.001).

### Statistical analyses

All analyses were conducted in R 4.4.2 (R Core Team, 2024). Linear mixed-effects models (LMMs) and generalised linear mixed-effects models (GLMMs) were fitted using the lme4 package (Bates et al., 2015). Treatment effects on endocrine and behavioural traits associated with parental investment were first examined, followed by offspring growth, telomere dynamics, and survival. Biologically relevant two-way interactions in all applicable models were initially included. We sequentially removed those with limited statistical support (*p* > 0.10), starting with the least supported interaction, while retaining the corresponding main effects. Model assumptions were assessed using the car (Fox & Weisberg, 2019) and DHARMa (Hartig, 2022) packages, as appropriate, including residual normality, homoscedasticity, simulated residual uniformity, and overdispersion where relevant.

### Parental endocrine and behavioural responses

Plasma prolactin concentrations were log-transformed and analysed using an LMM, with treatment, parental sex, breeding stage (i.e. incubation vs. chick rearing), and their two-way interactions as initial fixed effects. Pre-implantation prolactin concentrations (D0) and blood-sampling duration were included as covariates to account for initial inter-individual variation and potential time-of-sampling effects, respectively. Parent identity was included as a random effect to account for repeated measurements. Time spent at sea was analysed using the same model structure, but without covariates. Probability of chick care behaviour was analysed using a binomial GLMM, with the number of scans in which chick care was observed versus not observed specified as a two-column response using the *cbind()* function. Treatment, days since implantation, and their interaction were included as fixed effects, while parent and observer identities were included as random effects.

### Offspring growth and survival

Hatching success was analysed using a binomial generalised linear model (GLM), with treatment as a fixed effect. Chick mass and structural size were analysed separately using LMMs, with treatment, chick sex, chick age, and their two-way interactions as initial fixed effects, and chick identity as a random effect to account for repeated measurements. Changes in structural size with chick age was used to assess structural growth. For chick mass, exploratory post-hoc comparisons between treatment groups at each chick age were conducted using estimated marginal means (emmeans package; Lenth & Piaskowski, 2026), with Tukey’s method for pairwise comparisons. Percentage differences in chick mass relative to the Placebo group were calculated using model-estimated marginal means. Chick survival was assessed using Kaplan–Meier survival curves and Cox proportional hazards models (survival package; Therneau, 2020), with treatment and chick sex as explanatory variables. We measured survival time from hatching until death or fledging, treating fledging as a right-censoring event.

### Offspring telomere dynamics

Relative telomere length was analysed using a LMM with treatment, chick sex, chick age, and their two-way interactions as initial fixed effects. Chick identity and qPCR plate identity were included as random effects to account for repeated measurements and inter-plate variation, respectively.

## Results and discussion

A 9% decline in circulating parental prolactin concentrations was observed following corticosterone treatment (F_1_,_40_._84_ = 4.70, p = 0.036; Fig. 1A; Table S1), with no significant variation in this effect across breeding stages (F_1_,_86_._30_ = 0.83, p = 0.36; Fig. 1A; Table S1), suggesting that experimentally elevated corticosterone alters an endocrine pathway supporting parental investment, consistent with previous experimental studies in seabirds (Angelier et al., 2009; Spée et al., 2011). Interestingly, circulating corticosterone concentrations progressively declined after implantation, reaching levels comparable to those of Placebo parents by the second foraging trip (Cotton et al., 2026). This suggests that the reduction in prolactin may have persisted beyond the period of elevated corticosterone. A similar dissociation was reported in black-legged kittiwakes (Angelier et al., 2009). This prolonged reduction in prolactin may reflect lasting neuroendocrine changes (Sharp et al., 1998; Angelier & Chastel, 2009) or altered energetic condition (Cherel et al., 1994; Groscolas et al., 2008; Angelier et al., 2009). The latter would be consistent with the progressive decline of body condition observed in CORT-treated parents (Cotton et al., 2026), and previously observed under severe energetic constraints in king penguins (Groscolas et al., 2008). Despite reduced prolactin concentrations, parental chick care did not differ significantly between treatments, with predicted probabilities of 3.2% in Placebo and 2.6% in CORT-treated parents (χ²_1_ = 0.87, p = 0.36; Table S2). Chick care declined significantly with time since implantation (χ²_1_ = 17.90, p < 0.001), with no significant treatment × time interaction (χ²_1_ = 1.80, p = 0.18; Table S2).

**Fig 1.**
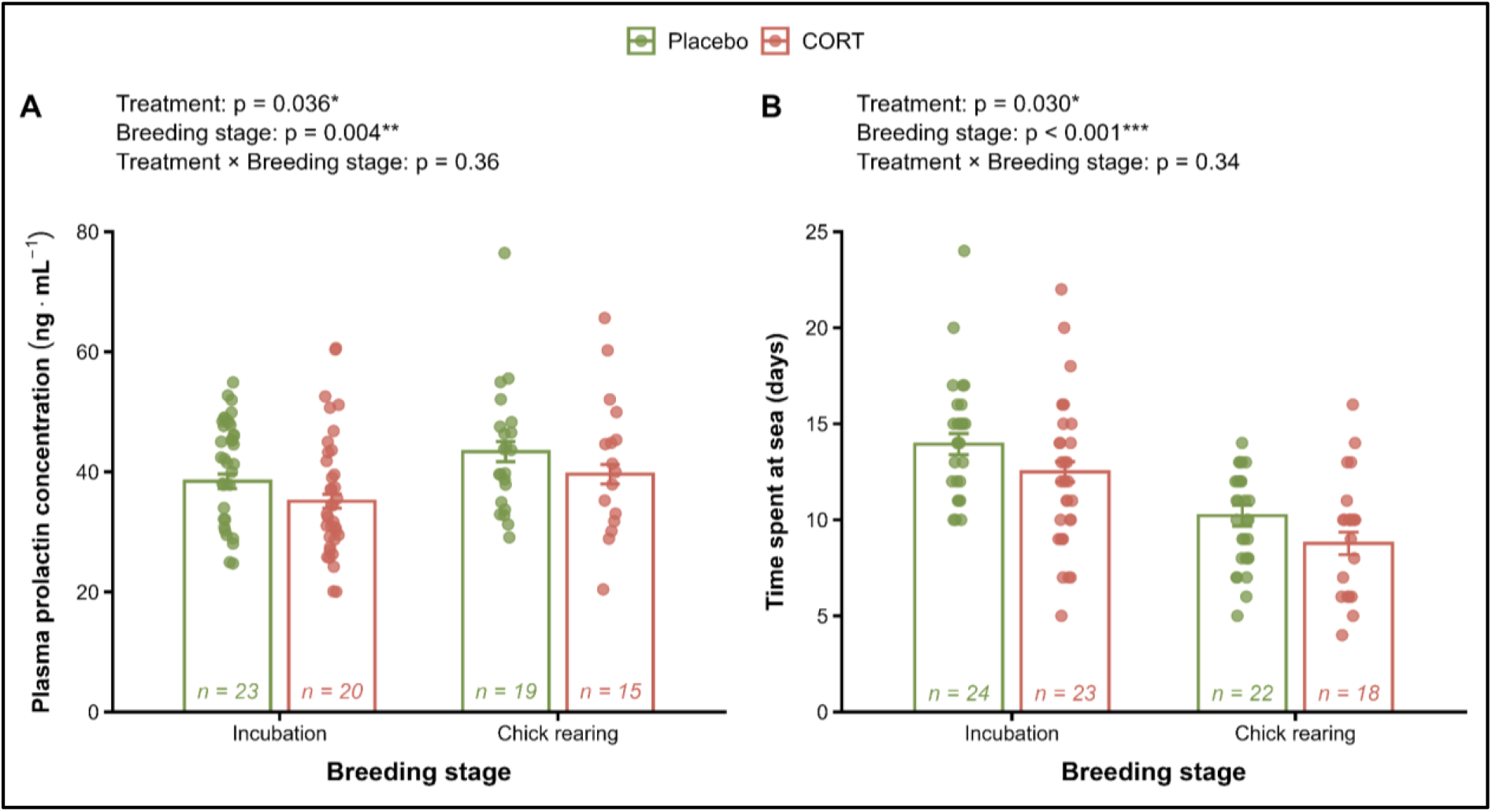
Effects of experimental corticosterone elevation in breeding king penguins on (A) prolactin hormone levels and (B) time spent at sea across breeding stages. Placebo and CORT-treated parents are shown in green and red, respectively. Points show raw individual values, while open bars and error bars represent estimated marginal means ± SE of linear mixed models. We analysed prolactin after log transformation, but present estimates on the original scale (ng·mL⁻¹).

In contrast, CORT-treated parents spent approximately 1.4 days less at sea per foraging trip than Placebo parents (treatment: F_1_,_4_₅ = 5.03, p = 0.030; Fig. 1B; Table S3), independently of breeding stage (treatment × breeding stage: F_1_,_8_₂._79_ = 0.92, p = 0.34). This shorter absence from the colony should have increased exposure to reproductive stimuli, such as visual and tactile cues from chicks, which usually stimulate prolactin secretion (Angelier & Chastel, 2009). Thus, reduced exposure to reproductive stimuli, as proposed in black-legged kittiwakes (Angelier et al., 2009), seems unlikely to explain the lower prolactin concentrations observed in CORT-treated parents. Shorter foraging trips are often associated with favourable environmental conditions and/or higher breeding success in seabirds (Uttley et al., 1994; Suryan et al., 2000; Thorne et al., 2015), including king penguins (Le Vaillant et al., 2016). However, shorter trips may reflect either faster resource acquisition or lower food intake. Experimental studies have shown that elevated corticosterone can modify foraging behaviour and energy expenditure in seabirds, with responses varying across species and environmental contexts (Angelier et al., 2008; Cottin et al., 2011; Crossin et al., 2012; Thierry et al., 2013a). Considering the progressive decline in parental condition (Cotton et al. 2026), the shorter foraging trips observed here in CORT-treated parents is unlikely to be associated with improved foraging performance.

Parental corticosterone treatment did not significantly affect hatching success (80.0% in CORT-treated pairs versus 91.7% in Placebo pairs; χ²_1_ = 1.40, p = 0.24). Chick mass showed no significant treatment × age interaction (F₅,_116_._96_ = 2.15, p = 0.065; Fig. 2A), although exploratory age-specific comparisons revealed that chicks raised by CORT-treated parents were 19.3% lighter at day 150 (estimated difference = −1.25 kg; p = 0.027), differences at other ages remained below 15% and were non-significant (all p ≥ 0.21; Fig. 2A; Table S4). Similarly, chick structural size showed neither a significant treatment effect (F_1_,_40_._68_ = 0.69, p = 0.41) nor a treatment × age interaction (F_4_,_79_._43_ = 1.22, p = 0.31; Fig. S2). This suggests broadly comparable growth patterns among surviving chicks despite parental corticosterone treatment. This maintenance of growth could reflect compensation by untreated partners or the selective disappearance of lighter or smaller chicks, potentially more vulnerable to predation (Descamps et al., 2005), although neither mechanism could be directly tested here. Parental corticosterone treatment significantly reduced chick survival (χ²_1_ = 3.88, p = 0.049; Fig. 2B, Table S6), with chicks raised by CORT-treated parents experiencing nearly twice the mortality hazard of those raised by Placebo parents (hazard ratio = 1.97, 95% CI = 1.01–3.86). A similar contrast between chick growth and survival was observed in Adélie penguins (*Pygoscelis adeliae*), with experimentally elevated parental corticosterone increasing chick mortality without significantly affecting mass among survivors (Thierry et al., 2013a). Survival curves appeared to diverge mainly after 35 days post-hatching, around 63.3 ± 8.7 days after parental implantation, when the experimentally induced elevation in parental corticosterone had already subsided (Cotton et al., 2026). This temporal mismatch between parental corticosterone exposure and offspring mortality suggests that the reproductive costs of parental stress may remain hidden during early development, only becoming apparent long after the initial physiological challenge has subsided.

**Fig. 2.**
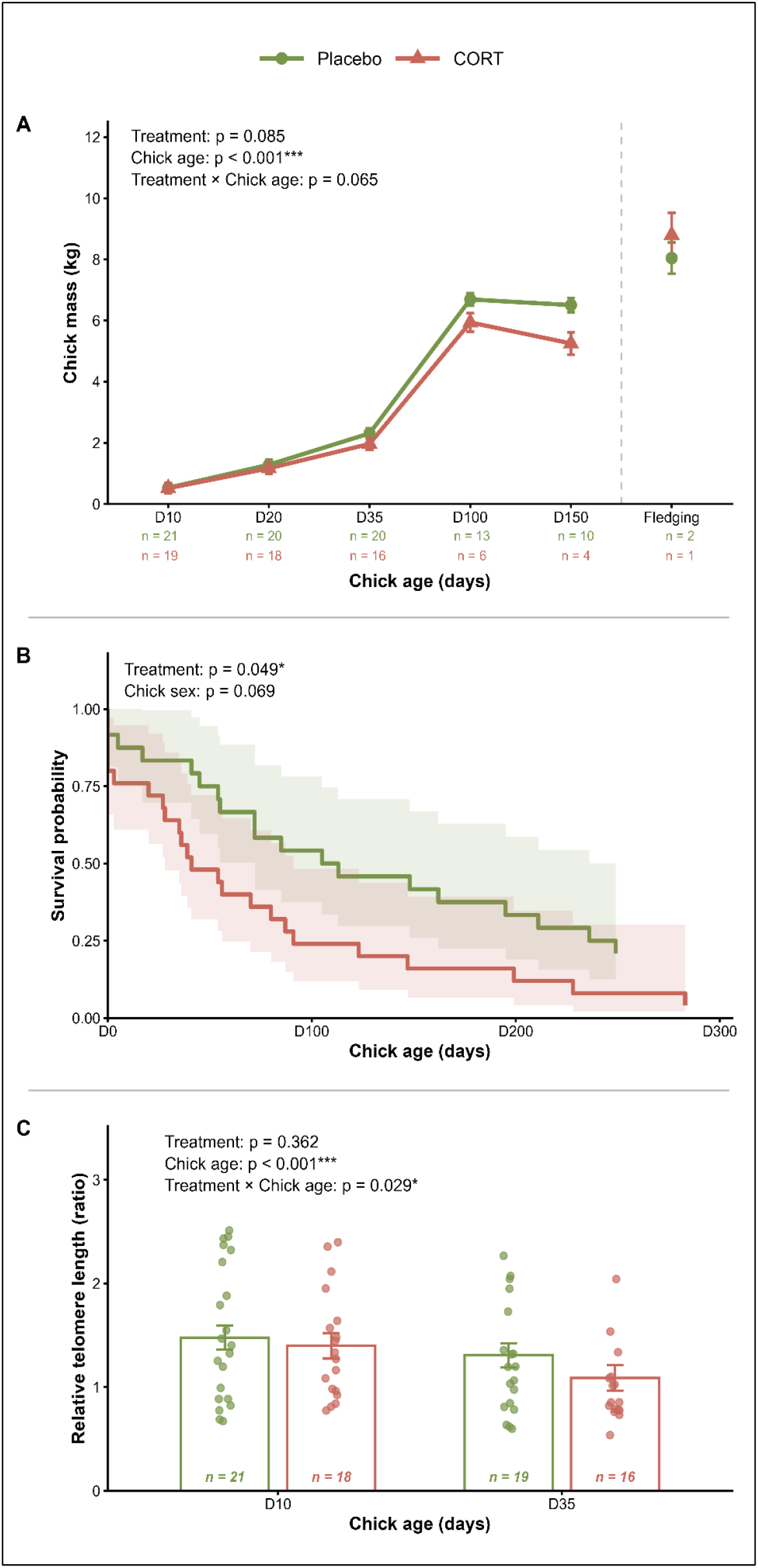
Effects of experimental parental corticosterone elevation on chick (A) mass, (B) survival and (C) relative telomere length. Offspring of Placebo and CORT-treated parents are shown in green and red, respectively. In A, symbols and error bars represent estimated marginal means ± SE, and fledging measurements are displayed separately from growth trajectories because age at fledging varied among individuals; in B, lines show Kaplan–Meier survival curves with 95% confidence intervals; and in C, open bars and error bars represent estimated marginal means ± SE, coloured points show individual values, and numbers indicate sample sizes.

Physiological differences between offspring of placebo and CORT-treated parents were already detectable before the apparent divergence in survival. Indeed, chick relative telomere length declined between 10 and 35 days post-hatching in both treatment groups, but this decline was significantly greater in chicks raised by CORT-treated parents (treatment × chick age: F_1_,_31_._39_ = 5.21, p = 0.029; Fig. 2C; Table S7). Telomere shortening was nearly twice as large in chicks from CORT-treated parents (ΔrTL = −0.31 ± 0.05, t_33_.₂ = −6.21, p < 0.001) as in Placebo (ΔrTL = −0.17 ± 0.04, t_31_._6_ = −4.14, p < 0.001). Early-life telomere dynamics have been associated with subsequent lifespan and lifetime reproductive success (Eastwood et al., 2019; Novelli et al., 2026), suggesting that these physiological changes may have implications beyond the period examined here. Challenging developmental conditions, including stress exposure and restricted resource availability, can accelerate telomere shortening (Reichert et al., 2015; Nettle et al., 2015). In king penguins, early-life telomeres are shorter under challenging colonial conditions (Stier et al., 2014), while their closer association with foster-parent rather than biological-parent telomere length highlights the importance of the rearing environment (Viblanc et al., 2020). Changes in parental provisioning or chick developmental conditions have likely contributed to accelerate telomere shortening in chicks from CORT-treated parents. The maintenance of chick mass and structural growth among surviving offspring, alongside greater telomere loss, raises the possibility that developmental costs were expressed at the cellular rather than morphological level. Such a pattern could reflect continued investment in growth at the expense of somatic maintenance under less favourable developmental conditions, as previously observed in last-hatched great tit nestlings (Stier et al., 2015). These mechanisms remain hypothetical, as we did not directly assess offspring energy allocation or compensatory parental provisioning. The effect of parental treatment on chick telomere length also differed between sexes (treatment × chick sex: F_1_,_3_₂._69_ = 7.70, p = 0.009), with shorter telomeres in male chicks raised by CORT-treated parents (Tukey post-hoc: males p = 0.011; females p = 0.21; Fig. S3). Sex-dependent effects of parental stress and glucocorticoid exposure on offspring telomere dynamics have previously been reported in other vertebrates (Marasco et al., 2019; Ensminger et al., 2021). Despite no detectable sex differences in chick mass or structural size (Tables S4 and S5), males may be more sensitive to developmental challenges, potentially owing to sex-specific sensitivity to oxidative stress (Giordano et al., 2015; López-Arrabé et al., 2016). Male king penguins are 15.5% heavier as adults (Pincemy et al., 2009), with size dimorphism emerging during development (Bordier et al., 2014), potentially increasing male energetic demands. Male chicks also showed an 87% higher estimated mortality hazard, although this was not statistically significant (p = 0.069; Table S6), so selective mortality of vulnerable males could partly obscure size dimorphism among survivors.

The physiological resilience of adult king penguins to prolonged corticosterone elevation (Cotton et al., 2026) may conceal important, delayed costs for their offspring. While early reproductive outcomes and overall chick growth were largely unaffected, accelerated telomere shortening and reduced survival revealed consequences that persisted or emerged well after parental corticosterone levels had returned to baseline. These responses, accompanied by reduced parental prolactin concentrations and shorter foraging trips, are consistent with changes in parental investment, potentially reflecting a compromise between self-maintenance and current reproduction. Thus, physiological resilience may not translate into reproductive resilience. Understanding how long-lived species balance survival and reproduction is important under environmental change. While recent studies have revealed how king penguins adjust their foraging strategies to environmental challenges (Lemonnier et al., 2025), linking parental foraging performance to the actual allocation of resources between self-maintenance and offspring provisioning remains a key challenge. Future studies that directly quantify food acquisition and delivery to chicks under experimentally elevated parental stress would help uncover the mechanisms linking parental decisions to offspring development and survival. To conclude, our study highlights the need to assess the consequences of environmental stressors over extended periods since deleterious effects may only emerge long after exposure. Such delayed effects are often difficult to assess but are of prime importance to better predict population dynamics in response to environmental changes.

## Acknowledgments

We thank the French Polar Institute (IPEV) and the Terres Australes et Antarctiques Françaises for financial and logistical support through the polar program #119 (ECONERGY). This work was conducted within the CNRS long-term Studies in Ecology and Evolution (SEE-Life) program. The CNRS Zone Atelier Antarctique et Terres Australes (ZATA) also provided additional financial support. We are also grateful to the staff of the Alfred Faure field station for their assistance and support during fieldwork, particularly Sylvain, Tom, and Pierre. AS received support from a Turku Collegium for Science and Medicine Fellowship, a Marie Skłodowska-Curie Postdoctoral Fellowship (#894963), and the IdEx Université de Strasbourg (*HotPenguin*). The French Polar Institute supported SA and LA.

## Data availability

The data used in this study is available as a supplementary file for reviewers, and will be publicly available upon acceptance in the CNRS InDoRES repository of the #119 penguin project (https://data.indores.fr/dataverse/ecoenergy119).

## Author contribution

Study design: AS, VAV, PBi. Funding acquisition: JPR, PBi, VAV, AS. Data collection in the field: AS, SA, TH, LA, and JPR. Data collection in the lab: ER, AS, AC & FA. Data analysis: AC, PBl, AS. Writing original draft: AC and AS. Writing review and editing: PBl, FA, VAV, JPR, PBi, SA, TH, LA, ER.

## Ethical note

The French Ethical Committee (APAFIS#16465-2018080111195526 v4) and the Terres Australes et Antarctiques Françaises (Arrêté TAAF A-2018-118) approved all procedures.

## Declaration of no competing interests

The authors declare no competing interests.

