## Supplementary figures and tables for "The cost of stressing the royal family: parental glucocorticoid exposure has delayed adverse effects on chick physiology and survival in king penguins"

### Supplementary materials

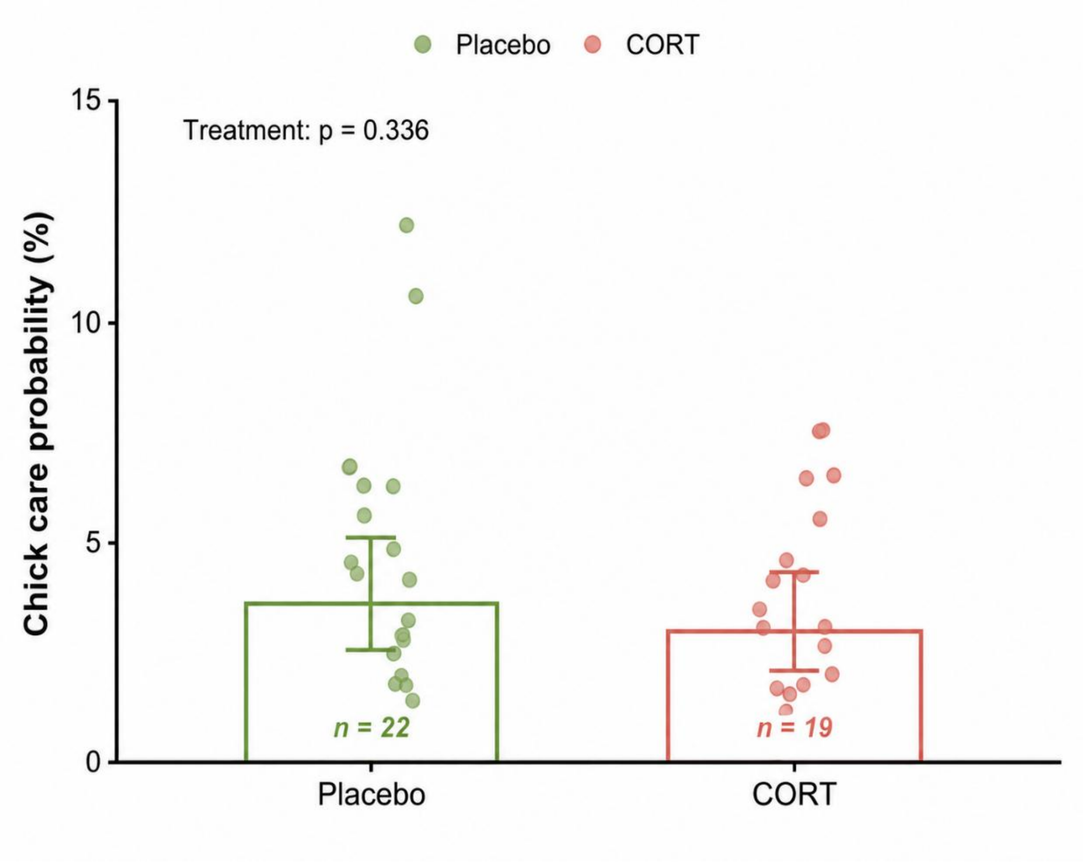


**Fig. S1. Effects of experimental parental corticosterone elevation on chick care probability in king penguins.** Placebo and CORT-treated parents are shown in green and red, respectively. Open bars represent estimated marginal probabilities of chick care, with error bars indicating 95% confidence intervals. Coloured points represent individual parent-level proportions of observations during which chick care was recorded, and numbers indicate sample sizes.

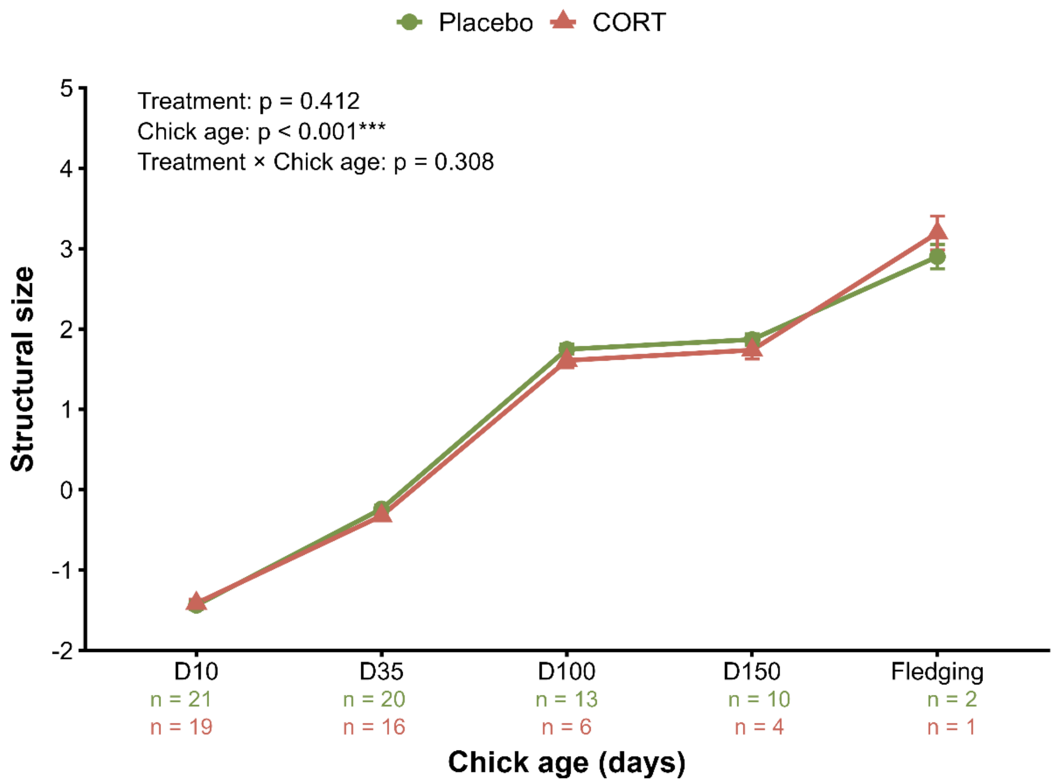


**Fig. S2. Effects of experimental parental corticosterone elevation on chick structural size.** Offspring of Placebo and CORT-treated parents are shown in green and red, respectively. Symbols and error bars represent estimated marginal means ± SE from the linear mixed model across chick ages. Lines connect estimated marginal means within each parental treatment, and sample sizes for each treatment and age are reported below the graph.

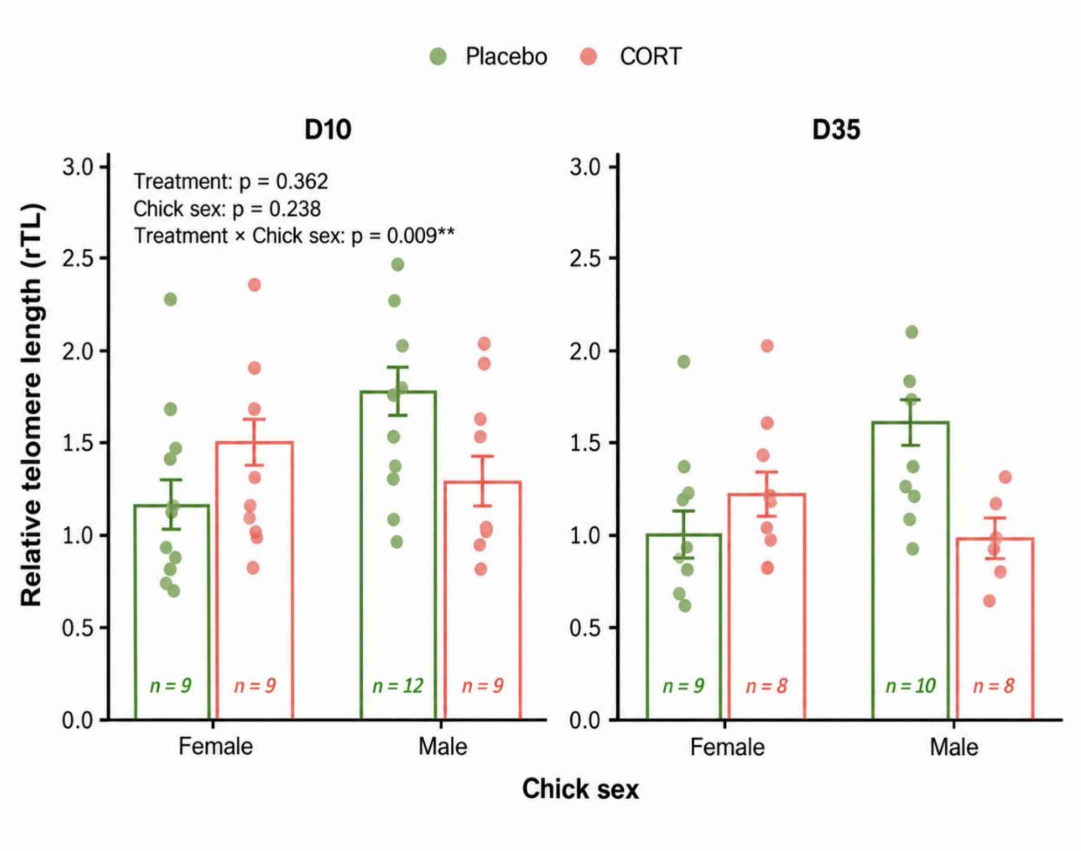


**Fig. S3. Sex-specific effects of experimental parental corticosterone elevation on chick relative telomere length at D10 and D35.** Offspring of Placebo and CORT-treated parents are shown in green and red, respectively. Female and male chicks are shown separately at D10 and D35. Open bars and error bars represent estimated marginal means ± SE from the linear mixed model; coloured points represent individual raw data; and numbers indicate sample sizes for each treatment, sex, and age.

Table S1. Results of the linear mixed-effects model on log-transformed prolactin levels (n = 111), with significant effects highlighted in bold. Non-significant interactions are reported below the horizontal line for completeness but were not retained in the final model presented above.

| **Random effects:** | | **Variance** | | | | | |
|---|---|---|---|---|---|---|---|
| Adults ID (N=45) | Intercept | 0.003 | | | | | |
| Residual | | 0.038 | | | | | |
| **Fixed effects:** | **df** | **F** | **p** | | **Estimate** | **Std. Error** | **t** |
| Intercept | | | | | 3.53 | 0.142 | 24.84 |
| **Treatment** | **1, 40.84** | **4.70** | **0.036** | **CORT** | **-0.09** | **0.042** | **-2.17** |
| **Parent sex** | **1, 41.54** | **24.89** | **<0.001** | **Male** | **-0.23** | **0.046** | **-4.99** |
| **Breeding stage** | **1, 85.28** | **8.74** | **0.004** | **Guard phase** | **0.12** | **0.041** | **2.96** |
| **Level at Day 0** | **1, 42.02** | **5.48** | **0.024** | | **0.008** | **0.003** | **2.34** |
| Blood sampling duration | 1, 103.70 | 2.39 | 0.125 | | -0.02 | 0.014 | -1.50 |
| Parent sex x Breeding stage | 1, 86.60 | 3.20 | 0.077 | Male x Guarding phase | | | |
| Breeding stage x Treatment | 1, 86.30 | 0.83 | 0.36 | Guarding phase x CORT | | | |

Table S2. Results of the generalised mixed-effects model on chick care (n = 325), with significant effects highlighted in bold. Non-significant interactions are reported below the horizontal line for completeness but were not retained in the final model presented above.

| **Random effects:** | | **Variance** | | | | | |
|---|---|---|---|---|---|---|---|
| Adults ID (N=41) | Intercept | 0.36 | | | | | |
| Observer (N=4) | Intercept | 0.28 | | | | | |
| **Fixed effects:** | **df** | **F** | **p** | | **Estimate** | **Std. Error** | **t** |
| Intercept | | | | | -2.33 | 0.40 | -5.78 |
| Treatment | 1 | 0.87 | 0.36 | CORT | -0.21 | 0.22 | -0.93 |
| **Days after implantation** | **1** | **17.90** | **<0.001** | | **-0.026** | **0.006** | **-4.23** |
| Treatment × Days after implantation | 1 | 1.80 | 0.18 | CORT × Days after implantation | | | |

Table S3. Results of the linear mixed-effects model on adult time spent at sea (n = 111), with significant effects highlighted in bold. Non-significant interactions are reported below the horizontal line for completeness but were not retained in the final model presented above.

| **Random effects:** | | **Variance** | | | | | |
|---|---|---|---|---|---|---|---|
| Adults ID (N=48) | Intercept | 0.81 | | | | | |
| Residual | | 9.35 | | | | | |
| **Fixed effects:** | **df** | **F** | **p** | | **Estimate** | **Std. Error** | **t** |
| Intercept | | | | | 13.67 | 0.67 | 20.53 |
| **Treatment** | **1, 45.00** | **5.03** | **0.030** | **CORT** | **-1.44** | **0.64** | **-2.24** |
| **Breeding stage** | **1, 82.21** | **39.43** | **<0.001** | **Guard phase** | **-3.72** | **0.59** | **-6.28** |
| Parent sex | 1, 52.21 | 0.67 | 0.42 | Male | 0.54 | 0.66 | 0.82 |
| Treatment x Breeding stage | 1, 82.79 | 0.92 | 0.34 | CORT x Guard phase | | | |
| Treatment x Parent sex | 1, 51.22 | 0.05 | 0.82 | CORT x Male parent | | | |

Table S4. Results of the linear mixed-effects model on chick mass (n = 150), with significant effects highlighted in bold. Non-significant interactions are reported below the horizontal line for completeness but were not retained in the final model presented above.

| **Random effects:** | | **Variance** | | | | | |
|---|---|---|---|---|---|---|---|
| Chicks ID (N=40) | Intercept | 0.097 | | | | | |
| Residual | | 0.483 | | | | | |
| **Fixed effects:** | **df** | **F** | **p** | | **Estimate** | **Std. Error** | **t** |
| Intercept | | | | | 0.756 | 0.17 | 4.54 |
| Treatment | 1, 43.98 | 3.11 | 0.085 | CORT | -0.27 | 0.15 | -1.76 |
| **Chick age** | **5, 121.30** | **292.44** | **<0.001** | **D20** | **0.71** | **0.16** | **4.47** |
| | | | | **D35** | **1.63** | **0.16** | **10.15** |
| | | | | **D100** | **5.90** | **0.20** | **29.65** |
| | | | | **D150** | **5.59** | **0.22** | **24.97** |
| | | | | **DFMU** | **7.76** | **0.44** | **17.83** |
| Chick sex | 1, 42.51 | 1.90 | 0.18 | Male | -0.21 | 0.15 | -1.38 |
| Treatment x Chick age | 5, 116.96 | 2.15 | 0.065 | CORT X D20 | | | |
| | | | | CORT x D35 | | | |
| | | | | CORT x D100 | | | |
| | | | | CORT x D150 | | | |
| | | | | CORT x DFMU | | | |
| Treatment x Chick sex | 1, 43.11 | 0.20 | 0.66 | | | | |

Table S5. Results of the linear mixed-effects model on chick structural size (n = 112), with significant effects highlighted in bold. Non-significant interactions are reported below the horizontal line for completeness but were not retained in the final model presented above.

| **Random effects:** | | **Variance** | | | | | |
|---|---|---|---|---|---|---|---|
| Chick ID (N = 40) | Intercept | 0.0130 | | | | | |
| Residual | | 0.0378 | | | | | |
| **Fixed effects:** | **df** | **F** | **p** | | **Estimate** | **Std. Error** | **t** |
| Intercept | | | | | -1.387 | 0.053 | -25.98 |
| Treatment | 1, 40.68 | 0.69 | 0.412 | CORT | -0.044 | 0.054 | -0.83 |
| **Chick age** | **4, 82.36** | **1199.30** | **<0.001** | **D35** | **1.150** | **0.045** | **25.57** |
| **Chick age** | | | | **D100** | **3.126** | **0.056** | **55.39** |
| **Chick age** | | | | **D150** | **3.252** | **0.064** | **51.09** |
| **Chick age** | | | | **Fledging** | **4.422** | **0.124** | **35.68** |
| Chick sex | 1, 39.04 | 0.52 | 0.477 | Male | −0.038 | 0.053 | −0.72 |
| Treatment x Chick age | 4, 79.43 | 1.22 | 0.308 | Placebo x D10 | | | |
| | | | | Placebo x D35 | | | |
| | | | | Placebo x D100 | | | |
| | | | | Placebo x D150 | | | |
| Treatment x Chick sex | 1, 39.85 | 0.05 | 0.832 | Placebo x Male chick | | | |

Table S6. Results of the Cox proportional hazards model on chick survival from hatching to fledging (n = 40 chicks, 35 deaths), with significant effects highlighted in bold. Hazard ratios (HR) and their 95% confidence intervals are presented, with Placebo-treated parents and female chicks as reference categories. Chi-square statistics and p-values are based on Type II likelihood-ratio tests.

| Fixed effects | df | $\chi^2$ | p | Level | Estimate | Std. Error | HR |
|---|---|---|---|---|---|---|---|
| **Treatment** | **1** | **3.88** | **0.049** | **CORT** | **0.679** | **0.343** | **1.97** |
| Chick sex | 1 | 3.31 | 0.069 | Male chick | 0.627 | 0.347 | 1.87 |
| Treatment x Chick sex | 1 | 0.287 | 0.592 | CORT x Male chick | | | |

Table S7. Results of the linear mixed-effects model on chick relative telomere length (n = 74), with significant effects highlighted in bold.

| **Random effects:** | | **Variance** | | | | | |
|---|---|---|---|---|---|---|---|
| Chick ID (N=39) | Intercept | 0.24 | | | | | |
| Plate ID (N=5) | Intercept | 0.004 | | | | | |
| Residual | | 0.016 | | | | | |
| **Fixed effects:** | **df** | **F** | **p** | | **Estimate** | **Std. Error** | **t** |
| Intercept | | | | | 1.16 | 0.17 | 6.79 |
| Treatment | 1, 31.80 | 0.85 | 0.36 | CORT | 0.37 | 0.24 | 1.56 |
| Chick sex | 1, 33.05 | 1.44 | 0.24 | Male | 0.64 | 0.22 | 2.90 |
| **Chick age** | **1, 31.57** | **61.16** | **<0.001** | **D35** | **-0.17** | **0.041** | **-4.18** |
| **Treatment x Chick sex** | **1, 32.69** | **7.70** | **0.009** | **CORT x Male chick** | **-0.89** | **0.32** | **-2.77** |
| **Treatment x Chick age** | **1, 31.39** | **5.21** | **0.029** | **CORT x D35** | **-0.14** | **0.061** | **-2.28** |
